# Two redundant ubiquitin-dependent pathways of BRCA1 localization to DNA damage sites

**DOI:** 10.1101/2021.07.21.452958

**Authors:** Alana Sherker, Natasha Chaudhary, Salomé Adam, Sylvie M. Noordermeer, Amélie Fradet-Turcotte, Daniel Durocher

## Abstract

The tumor suppressor BRCA1 accumulates at sites of DNA damage in a ubiquitin-dependent manner. In this work, we revisit the role of the ubiquitin-binding protein RAP80 in BRCA1 recruitment to damaged chromatin. We found that RAP80, or the phosphopeptide-binding residues in the BRCA1 BRCT domains, act redundantly with the BRCA1 RING domain to promote BRCA1 recruitment to DNA double-strand break sites. We show that that RNF8 E3 ubiquitin ligase acts upstream of both the RAP80- and RING-dependent activities whereas RNF168 acts uniquely upstream of the RING domain. The function of the RING domain in BRCA1 recruitment is not solely linked to its role in mediating an interaction with BARD1 since RING mutations that do not impact BARD1 interaction, such as the E2-binding deficient Ile26Ala (I26A) mutation, produce a BRCA1 protein unable to accumulate at DNA damage sites in the absence of RAP80. Cells that combine the BRCA1 I26A mutation and mutations that disable the RAP80-BRCA1 interaction are deficient in RAD51 filament formation and are hypersensitive to poly (ADP-ribose) polymerase inhibition. Our results suggest that in the absence of RAP80, the BRCA1 E3 ligase activity is necessary for the recognition of unmethylated histone H4 Lys20 and histone H2A Lys13/Lys15 ubiquitylation by BARD1 although we cannot rule out the possibility that the RING- E2 complex itself may facilitate ubiquitylated nucleosome recognition in other ways. Finally, given that tumors expressing RING-less BRCA1 isoforms readily acquire resistance to therapy, this work suggests that targeting RAP80, or its interaction with BRCA1, could represent a novel strategy for restoring sensitivity of such tumors to DNA damaging agents.

## Introduction

BRCA1 is encoded by the first familial breast and ovarian cancer tumor suppressor gene identified (Futreal et al., 1994; Miki et al., 1994). The mechanism by which BRCA1 suppresses oncogenesis is most likely linked to its function in activating DNA repair by homologous recombination (HR) (Bhattacharyya et al., 2000; Moynahan et al., 1999) although other mechanisms have also been proposed (Tarsounas and Sung, 2020). BRCA1 localizes to sites of DNA damage (Paull et al., 2000; Scully et al., 1997a; Scully et al., 1997b), implying that BRCA1 acts to promote DNA repair directly at DNA lesions but this link has yet to be formally established. BRCA1 is a modular protein of 1863 amino acid residues with a RING finger domain located at the N-terminus and a coiled-coil region as well as two tandem BRCT domains at the C-terminus (Figure 1a). BRCA1 forms an obligatory heterodimer with the BRCA1-associated RING domain protein (BARD1) through an interaction via their respective RING finger domains (Figure 1a). This interaction contributes to the stability of both proteins and confers E3 ubiquitin ligase activity to the BRCA1- BARD1 complex towards the C-terminus of histone H2A, specifically the K125/K127/K129 residues (Becker et al., 2020; Boulton, 2006; Densham et al., 2016; Kalb et al., 2014; Nakamura et al., 2019; Polanowska et al., 2006; Wu et al., 1996).

**Figure 1.**
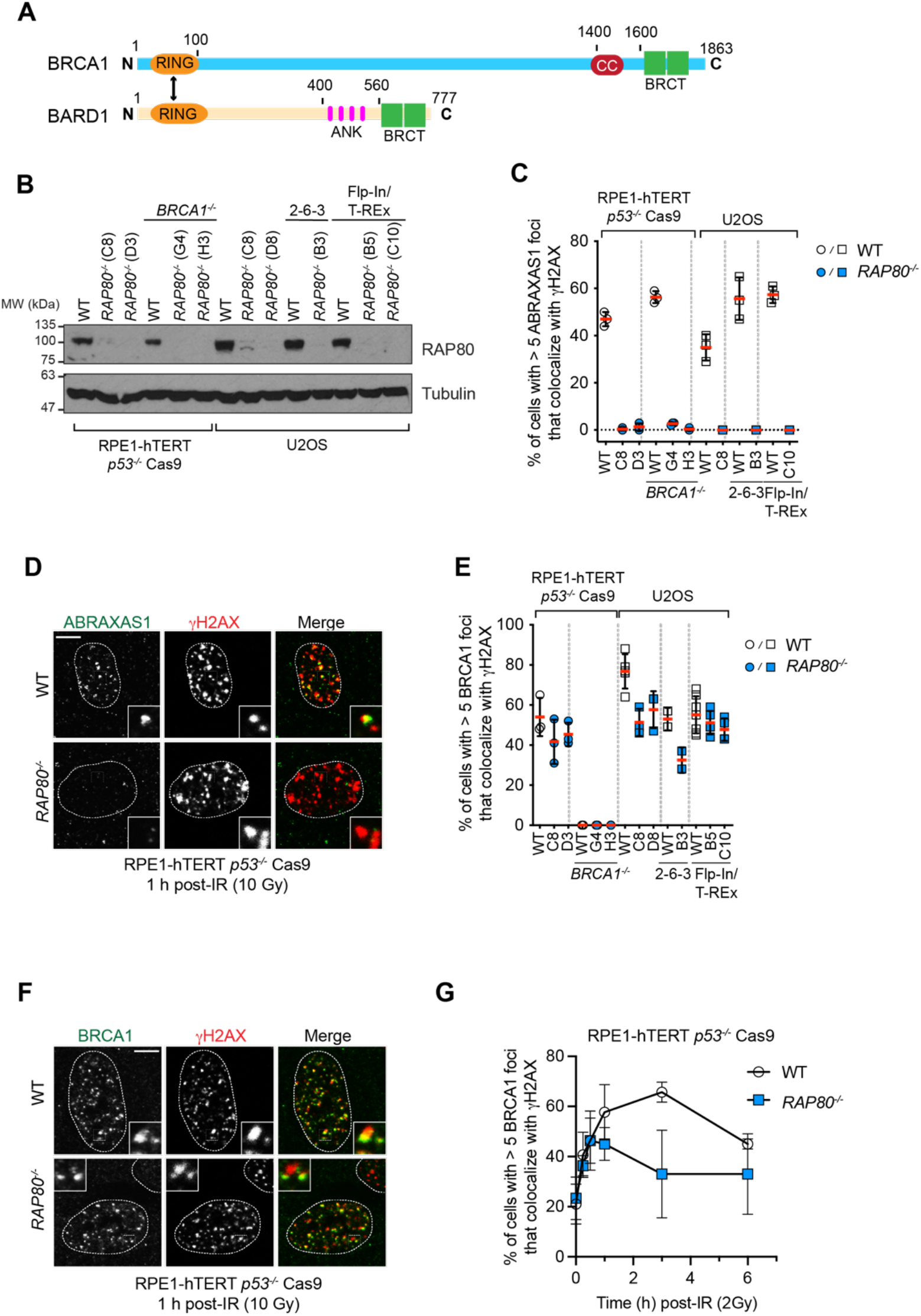
RAP80 is not necessary for BRCA1 recruitment to DSB sites. (A) Schematic representation of BRCA1 and BARD1. Highlighted are RING domains (orange), PALB2- interacting coiled-coil region (CC; red), BRCA1 C-terminal domains (BRCT; green) and ankyrin repeats (ANK; pink). (B) Immunoblotting of whole cell extracts obtained from parental and *RAP80^-/-^* clones in the RPE-1 hTERT *p53^-/-^* Cas9, RPE-1 hTERT *p53^-/-^ BRCA1^-/-^* Cas9, U2OS, U2OS-2-6-3 and U2OS Flp-In/TREx cell lines with RAP80 antibodies. Tubulin is used as loading control. See Figure S1 for further details on the gene editing strategy used to knockout *RAP80*. (C, D) The indicated parental and *RAP80^-/-^* cell lines were processed for immunofluorescence 1 h post- irradiation (10 Gy) and stained with antibodies against ABRAXAS1 and γH2AX. Shown in (C) is the percentage of cells with > 5 ABRAXAS1 foci that colocalize with γH2AX. A minimum of 100 cells per replicate were analyzed and the bars represent the mean ± S.D. (n=3). Representative micrographs of RPE1-hTERT *p53^-/-^* Cas9 wild type (WT) and *RAP90^-/-^* cells are shown in (D). (E, F) The indicated cell lines were processed for immunofluorescence 1 h post-irradiation (10 Gy) and stained with antibodies against BRCA1 and γH2AX. Shown in (E) is the percentage of cells with > 5 BRCA1 foci that colocalize with γH2AX. A minimum of 100 cells per replicate were analyzed and the bars represent the mean ± S.D. (n=3). Representative microgaphs of RPE-1 hTERT *p53^-/-^* Cas9 wild type (WT) and *RAP80^-/-^* are shown in (F). (G) RPE-1 hTERT *p53^-/-^* Cas9 parental (WT) and *RAP80^-/-^* cells were untreated (t=0 h) or irradiated with a 10 Gy dose. Samples were collected at the indicated time points, processed for immunofluorescence and stained with antibodies against BRCA1 and γH2AX. Shown is the percentage of cells with > 5 BRCA1 foci. A minimum of 100 cells per replicate were analyzed and the bars represent the mean ± S.D. (n=3). All scale bars are 5 μm.

BRCA1 accumulates on the chromatin surrounding DNA damage sites in a manner that depends on histone H2AX phosphorylation and the RNF8- and RNF168-catalyzed histone ubiquitylation cascade (Celeste et al., 2003; Doil et al., 2009; Huen et al., 2007; Kolas et al., 2007; Mailand et al., 2007; Sobhian et al., 2007; Stewart et al., 2009). BRCA1 does not contain any recognizable ubiquitin-binding domain, but interacts with BRCA1-A, a large ubiquitin-binding complex formed by the ABRAXAS, RAP80, BABAM1, BABAM2 and BRCC3 proteins (Kim et al., 2007b; Liu et al., 2007; Wang et al., 2007) (Kyrieleis et al., 2016; Rabl et al., 2019; Shao et al., 2009). BRCA1 binds to BRCA1-A via its tandem BRCT domains that recognize phosphorylated ABRAXAS1 (also known as Abraxas or FAM175A). Within BRCA1-A, the RAP80 subunit (also known as UIMC1) has high affinity for the Lys63-linked ubiquitin (UbK63) chains produced by RNF8 and RNF168; thereby providing a means for BRCA1 recruitment to DNA lesions (Hu et al., 2012; Kim et al., 2007a; Rabl et al., 2019; Sims and Cohen, 2009; Sobhian et al., 2007; Walters and Chen, 2009).

While this model of BRCA1 recruitment is attractive, loss of the BRCA1-A complex results in increased DNA end-resection and higher levels of HR detectable as gene conversion (Coleman and Greenberg, 2011; Dever et al., 2011; Hu et al., 2011), which is in contrast to the loss of end-resection and HR activity seen in BRCA1-deficient cells (Cruz-Garcia et al., 2014; Schlegel et al., 2006; Stark et al., 2004). These observations suggest either that BRCA1 localization to DSB sites is irrelevant for its function during HR or that there are elements of BRCA1 localization to DSB sites that remain unresolved. In support of the latter possibility, RNA interference studies showed that RAP80 was dispensable for the initial BRCA1 localization at DNA damage sites but was rather proposed to be involved in the maintenance of BRCA1 on damaged DNA (Hu et al., 2011). This work implied that other mechanisms of BRCA1 recruitment to ubiquitylated chromatin must exist.

In an effort to develop a better understanding of the mechanisms of BRCA1 recruitment to DNA damage sites, we revisited the contribution of RAP80 to BRCA1 localization to DSB sites using genetic knockouts of *RAP80* in multiple cell backgrounds. We found that *RAP80^-/-^* cells have robust BRCA1 localization to DSB sites and uncovered that this was due to near-complete redundancy with a DSB site-targeting activity that located in the BRCA1 RING finger domain. Mutations that alter BRCA1 RING function without impairing its BARD1 interaction (such as the E2 binding-deficient I26A mutation) causes complete loss of BRCA1 localization to DNA damage sites and greatly impair HR in the absence of RAP80 or in the presence of mutations that disable the BRCA1/BRCA1-A interaction. We conclude that DNA damage localization of BRCA1 is essential for its function during HR and that it is dependent on two redundant activities mediated by the BRCA1-A complex and the BRCA1 RING domain. We finally speculate that the BRCA1 E3 ligase activity may play an important role in endowing recognition of RNF168-catalyzed H2A Lys13/15 ubiquitylation (H2A-K13/K15ub) by the BARD1 protein (Becker et al., 2020).

## Results

To better understand the contribution of RNF168-dependent ubiquitylation to BRCA1 accumulation to DSB sites, we generated *RAP80* (*UIMC1*) knockouts by gene editing in multiple RPE1-hTERT *p53^-/-^* Cas9 (RPE1) and U-2-OS (U2OS) cell backgrounds (Figure 1b and Figure S1a). As expected, analysis of gene conversion using the traffic light reporter assay in RPE1 cells showed that loss of RAP80 causes an increase, rather than a decrease, in HR (Figure S1b). Furthermore, RAP80 inactivation also demonstrated a complete loss of ABRAXAS1 localization to ionizing radiation (IR)-induced foci marked by γH2AX in multiple RPE1- and U2OS-derived clones (Figure 1cd). These results indicated that these cell lines recapitulate known phenotypes associated with RAP80 inactivation.

These 7 independent *RAP80^-/-^* cell lines displayed near-normal recruitment of BRCA1 to DSB sites as measured by IR-induced focus formation (Figure 1ef). The BRCA1 foci were completely lost in *BRCA1^-/-^* cells (Figure 1ef) indicating that the immunostaining was specific for the BRCA1 protein. We next examined BRCA1 IR-induced focus formation and retention over time, fixing cells from 15 min to 6 h post-irradiation. We detected a defect in the maintenance of BRCA1 foci from 1 h onwards in the *RAP80^-/-^* cell line (Figure 1g and Figure S1c), consistent with the phenotypes previously described using short interfering (si) RNA-mediated depletion of RAP80 (Hu et al., 2011). We therefore conclude that RAP80, and by inference the BRCA1-A complex, is not necessary on its own for the initial recruitment of BRCA1 to DSB sites.

RAP80 interacts specifically with UbK63 chains. While both RNF8 and RNF168 participate in the formation of UbK63 chains, mounting evidence suggests that RNF8 is the main source of UbK63 at DNA damage sites by ubiquitylating histone H1 (Thorslund et al., 2015). We therefore tested whether there was a differential contribution for RNF8 or RNF168 towards BRCA1 recruitment in cell lines that were proficient or deficient in RAP80. We observed that RNF8 depletion by effective siRNAs (Fradet-Turcotte et al., 2013) led to a near-complete loss of BRCA1 recruitment to IR-induced foci, while depletion of RNF168 led, in comparison, to an incomplete decrease (Figure 2a,b). The residual BRCA1 recruitment to DSB sites observed in RNF168-depleted cells was dependent on RAP80 since depletion of RNF168 in *RAP80*^-/-^ cells led to a complete loss of BRCA1 IR-induced foci (Figure 2a,b). To rule out the possibility that these results were an artefact of siRNA-mediated depletion, we examined BRCA1 recruitment in *RNF8*^-/-^ and *RNF168*^-/-^ cell lines generated in RPE1-hTERT *p53^-/-^* Cas9 cells (Figure S1d). As with siRNA depleted cells, we observed in *RNF8*^-/-^ cells a loss of RNF8 caused complete loss of BRCA1 accumulation into IR-induced foci compared to a partial reduction of BRCA1 recruitment in the *RNF168* knockout cells (Figure 2c,d). Depletion of RAP80 in *RNF168*^-/-^ cells abolished the residual recruitment of BRCA1 to DNA damage sites (Figure 2c,d). Examination of localization of RAP80 to IR-induced foci in the *RNF8^-/-^* and *RNF168^-/-^* cells showed that BRCA1-A localization was completely dependent on RNF8 but only partially dependent on RNF168 (Figure S1e,f). These results indicate that RAP80 may promote a mode of BRCA1 recruitment to DNA damage sites that is largely dependent on RNF8 and that acts in parallel to a second mode of recruitment that is dependent on RNF168- mediated ubiquitylation of histone H2A Lys13/Lys15 residues.

**Figure 2.**
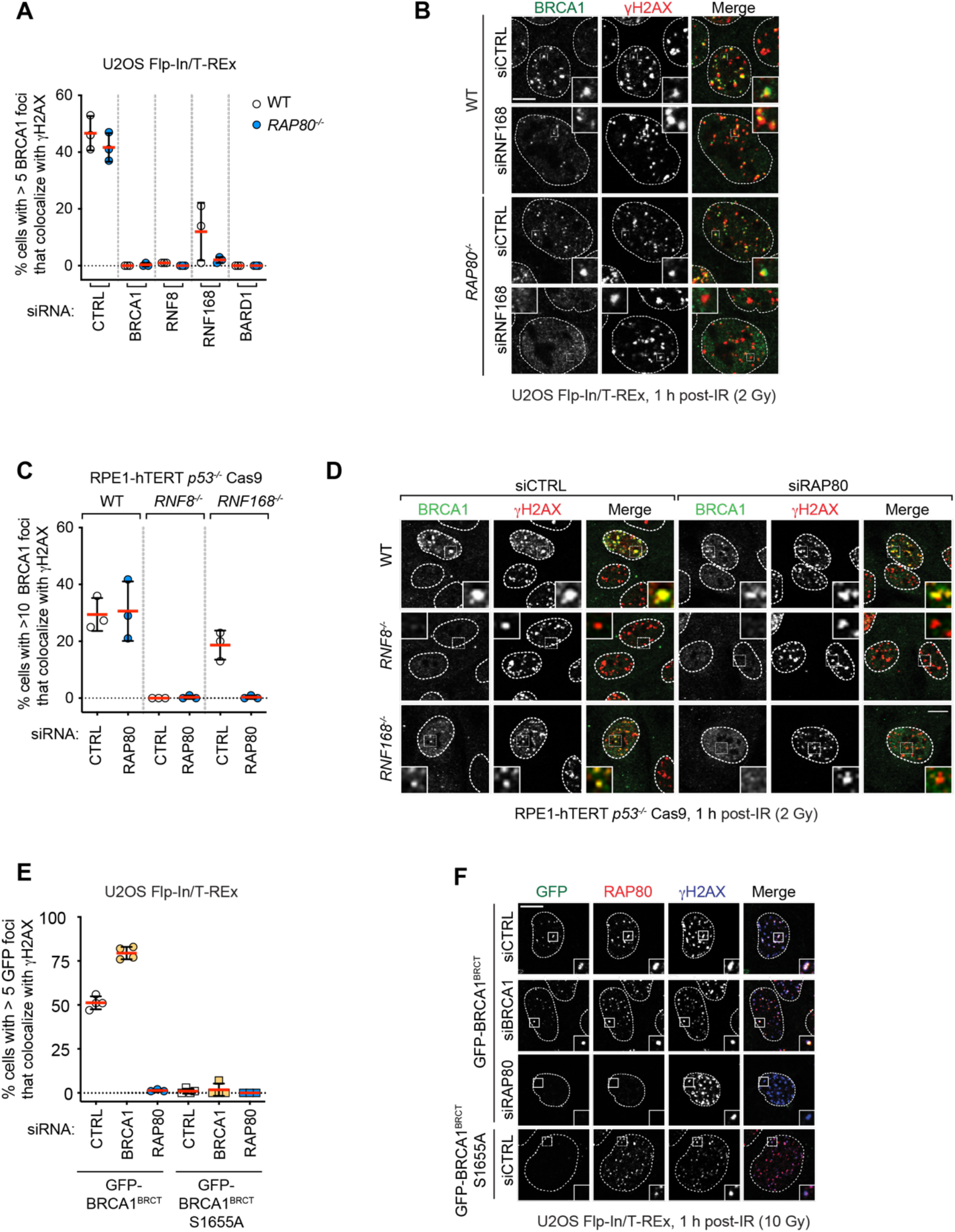
RAP80-independent BRCA1 recruitment to DSB sites is dependent on RNF8 and RNF168. (A, B) U2OS Flp-In/T-Rex parental (WT) and *RAP80^-/-^* cells were transfected with siRNA pools targeting *BRCA1*, *RNF8*, *RNF168* or *BARD1* or with a non-targeting control siRNA (CTRL). 48 h post-transfection cells were irradiated (2 Gy) and processed for immunofluorescence 1 h post-IR treatment using antibodies against BRCA1 and γH2AX. Quantitation of the percentage of cells with > 5 BRCA1 foci that colocalize with γH2AX is shown in (A). A minimum of 100 cells per replicate were analyzed and the bars represent the mean ± S.D. (n=3). Representative micrographs are shown in (B). (C, D) Parental (WT) RPE1-hTERT *p53^-/-^* Cas9, *RNF8^-/-^*, and *RNF168^-/-^* cells were treated with either a non-targeting siRNA pool (CTRL) or a pool targeting *RAP80*. 48 h post-transfection cells were irradiated (2 Gy) and processed for immunofluorescence 1 h post-IR treatment using antibodies against BRCA1 and γH2AX. Quantitation of the percentage of cells with > 10 BRCA1 foci that colocalize with γH2AX is shown in (C). A minimum of 100 cells per replicate were analyzed and the bars represent the mean ± S.D. (n=3). Representative micrographs are shown in (D). (E, F) U2OS Flp-In/T-REx cells with integrated transgenes encoding GFP-BRCA1^BRCT^ or -BRCA1^BRCT^ S1655A were transfected with non-targeting siRNA (CTRL) or siRNA targeting *RAP80* or *BRCA1*. Following doxycycline treatment to induce transgene expression (5 μg/mL, 24 h), cells were irradiated (10 Gy) and processed for immunofluorescence 1 h post-IR for GFP and antibodies against RAP80 and γH2AX. Shown in (E) is the quantitation of a minimum of 100 cells per replicate where the bars represent the mean ± S.D (n= 4). Representative micrographs are shown in (F). All scale bars are 5 μm.

To further dissect how BRCA1 may be recruited to DNA damage via RAP80-dependent and - independent pathways, we examined how a truncated protein comprised of the isolated tandem BRCT domains of BRCA1 (amino acid residues 1582-1863; BRCA1^BRCT^) is recruited to DNA damage sites. We observed that, contrary to the observed results for full-length BRCA1 recruitment, localization of BRCA1^BRCT^ into IR-induced foci was strictly dependent on RAP80 and the ABRAXAS1-interacting S1655 residue in the BRCT domains (Figure 2e,f). These results hinted that the putative second and RAP80-independent mode of recruitment of BRCA1 to DNA lesions is carried out by a BRCA1 region other than the tandem BRCT domains. In order to map this additional recruitment domain, we generated stable U2OS Flp-In/T-Rex cell lines that express various siRNA-resistant transgenes producing GFP-tagged BRCA1 and variants. Consistent with the results above, we observed that deletion of the BRCT domains or introduction of the S1655A phosphopeptide-binding mutant in the context of full-length BRCA1 maintain the ability of BRCA1 to form IR-induced foci (Figure 3a,b). Furthermore, the variant BRCA1 1-1362, containing a C-terminal deletion of both BRCT and the PALB2-interacting coiled coil regions, also formed robust IR-induced foci in U2OS cells (Figure 3a,b). However, to our surprise, expression of a protein consisting essentially solely of the RING finger domain (BRCA1^RING^ i.e. BRCA1 1-110) also localized to DNA damage sites independently of RAP80 (Figure 3a,b) with similar efficiency to the full-length protein when focus intensity was measured (Figure S2a). These results suggest that the RING domain may be responsible for an activity that recruits BRCA1 to DNA damage sites redundantly with RAP80.

**Figure 3.**
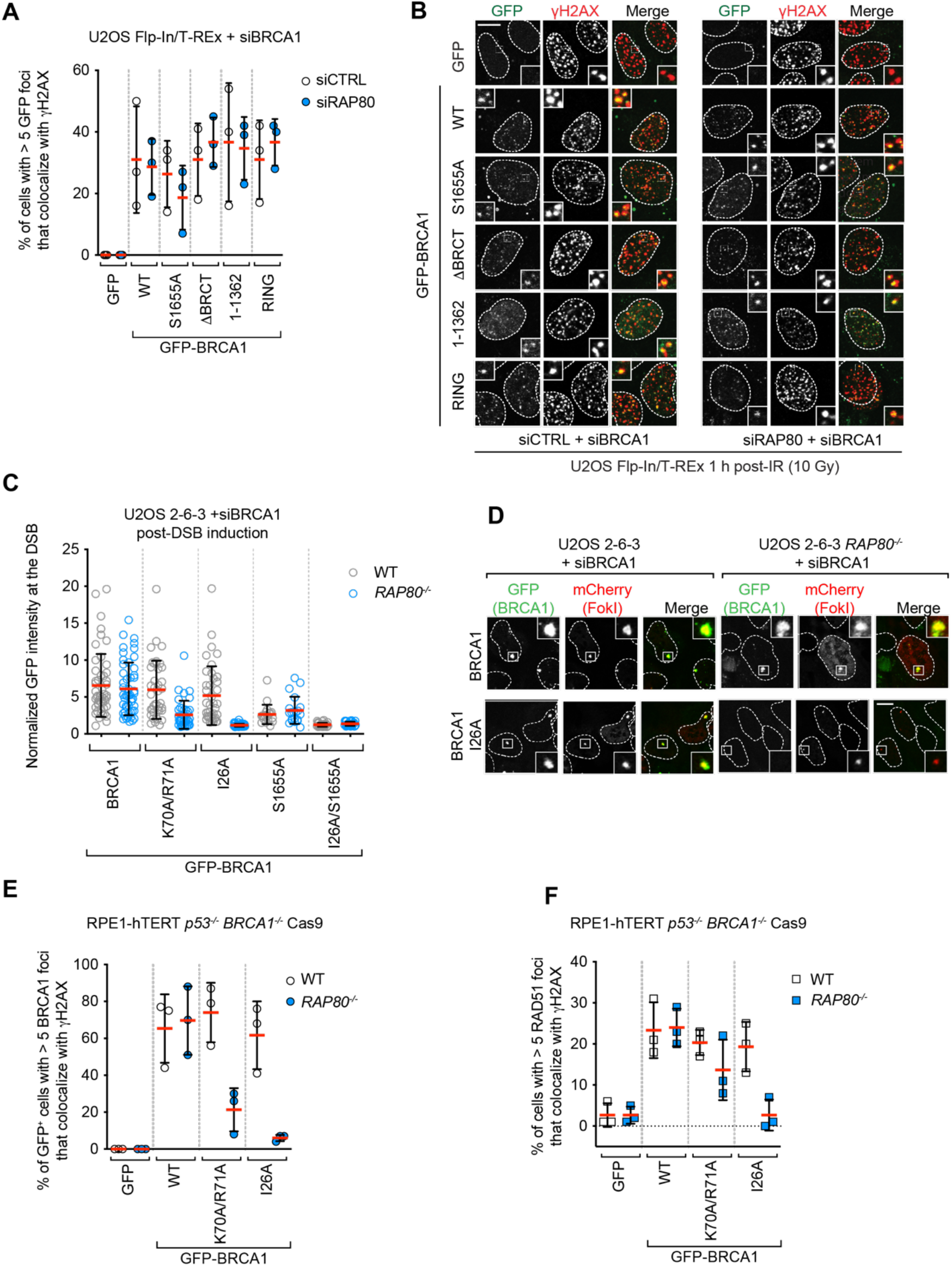
The RING domain participates in BRCA1 recruitment to DNA damage sites. (A,B) U2OS Flp-In/T-REx cells stably integrated with the indicated transgenes were treated with doxycycline (5 μg/mL, 36 h) to induce protein expression and transfected with an siRNA targeting *BRCA1* and also either non-targeting siRNA (CTRL) or siRNAs targeting *RAP80*. 1 h post-irradiation (10 Gy) cells were processed for immunofluorescence using GFP (to detect the BRCA1 fusions) and antibodies against γH2AX. Shown in (A) is the quantitation of a minimum of 100 cells per replicate where the bars represent the mean ± S.D (n= 3). Representative micrographs are shown in (B). (C, D) U2OS 2-6-3 parental (WT) or *RAP80^-/-^* cell lines were transfected with siRNAs targeting *BRCA1* followed by nucleofection of the indicated GFP fusion proteins. 48 h post-nucleofection, mCherry-LacR-FokI expression was induced for 5 h prior to being processed for fluorescence microscopy for GFP and mCherry. Shown in (C) is the quantitation of GFP fluorescence at the mCherry focus where the bars represent the mean ± S.D (n= 3). Representative micrographs for the BRCA1 and BRCA1-I26A conditions are shown in (D). Additional micrographs for the other conditions are in Figure S2c. (E) RPE-1 hTERT *p53^-/-^ BRCA1^-/-^* Cas9 cells (WT) or their isogenic *RAP80^-/-^* counterparts expressing the indicated GFP fusion proteins were processed 1 h post-irradiation (10 Gy) for immunofluorescence using GFP and antibodies against BRCA1 and γH2AX. A minimum of 100 cells per replicate were analyzed and the bars represent the mean ± S.D (n= 3). Representative micrographs are shown in Figure S3b. (F) RPE- 1 hTERT *p53^-/-^ BRCA1^-/-^* Cas9 cells or their isogenic *RAP80^-/-^* counterparts expressing the indicated GFP fusion proteins were processed 1 h post-irradiation (10 Gy) for immunofluorescence using GFP and antibodies against RAD51 and γH2AX. A minimum of 100 cells per replicate were analyzed and the bars represent the mean ± S.D (n= 3). Representative micrographs are shown in Figure S3c. Scale bars are 5 μm in (B) and 10 μm in (D).

Mutations or loss of the RING domain in BRCA1 impairs its association with BARD1 and leads to BRCA1 destabilization (Fabbro et al., 2002; Hashizume et al., 2001; Joukov et al., 2001), complicating the analysis of the contribution of the RING domain in BRCA1 recruitment. We therefore explored whether we could identify mutations in the BRCA1 RING domain that impair DNA damage localization while maintaining stability. We selected two mutations: BRCA1-I26A, which disrupts the interaction between the RING and E2 conjugating enzymes such as UbcH5c (Brzovic et al., 2003; Christensen et al., 2007) and BRCA1 K70A/R71A that disrupts the interaction between BRCA1 and the nucleosome acidic patch (McGinty et al., 2014; Witus et al., 2021). These two mutants, along with wild type BRCA1, were expressed as fusions to GFP from siRNA-resistant transgenes in U2OS 2-6-5 cell lines (parental and *RAP80^-/-^*) (Figure S2b). The U2OS 2-6-5 cell line contains an inducible mCherry-LacR-FokI fusion protein that can induce clustered DSBs at an integrated LacO array, which allows for facile quantitation of recruitment to DSB sites (Shanbhag et al., 2010). Upon depletion of BRCA1 by siRNA, FokI expression was induced and GFP fusion protein recruitment to mCherry-marked DSBs was assessed. We observed that the two BRCA1 RING mutants accumulated at DSB sites as efficiently as wild type BRCA1 in RAP80-proficient cells but had greatly impaired recruitment to FokI-induced breaks in *RAP80^- /-^* cells, with BRCA1 I26A being the most defective (Figure 3c,d and Figure S2c). These results suggested that BRCA1 recruitment to DSB sites is the result of a collaboration between the RING domain and the BRCT-dependent interaction with BRCA1-A. To test this idea, we also introduced the BRCA1-S1655A mutation alone or in combination with I26A. The S1655A mutant showed reduced but RAP80-independent recruitment to FokI-induced DSBs that was completely abolished by the I26A mutation (Figure 3c,d).

To further test the collaboration between RAP80 and the BRCA1 RING domain in an orthogonal system, we used gene editing to create a RAP80 knockout (*RAP80^-/-^*) in an RPE1-hTERT *BRCA1^- /-^ p53^-/-^* Cas9 cell line (Figure S3a). This cell line allowed us to assess BRCA1 IR-induced focus formation and its dependence on RAP80 in cell lines expressing BRCA1 variants. As observed with the FokI system, we found that the BRCA1 I26A protein form IR-induced foci but does so in a strictly RAP80-dependent manner (Figure 3e and Figure S3b). The accumulation of BRCA1 K70A/R71A at DSB sites was also largely dependent on RAP80 but to a lesser extent than for BRCA1 I26A (Figure 3e and Figure S3b). Together, these results indicate that both the nucleosome-RING and E2-RING interactions participate in the recruitment of BRCA1 to DSB sites in parallel to the BRCA1-A-dependent recognition of UbK63 chains by RAP80.

The identification of conditions where BRCA1 recruitment to DSB sites is severely impaired allowed us to ask whether BRCA1 accumulation into IR-induced foci correlates with DNA repair activity by BRCA1. We first assessed RAD51 IR-induced focus formation in the RPE1-hTERT cell lines described above as a proxy for HR activity. Mirroring BRCA1 recruitment, we found that RAD51 focus formation was near-normal in the RAP80-proficient cell lines expressing the BRCA1 K70A/R71A and I26A mutants but was impaired in the *RAP80*^-/-^ cell lines, with BRCA1- 26A showing reduction of RAD51 IR-induced foci to the levels of the *BRCA1^-/-^* cells expressing only GFP and (Figure 3f and Figure S3c). These data suggest that RING mutants of BRCA1 rely on their interaction with BRCA1-A for HR activity.

The RPE-hTERT *BRCA1^-/-^* cell lines above did not maintain homogenous expression of the BRCA1 variants long enough to allow an assessment of HR activity using assays such as resistance to PARP inhibition. We therefore used MDA-MB-436 cells, which harbor a hemizygous *BRCA1 5396+1G >A* mutation that causes complete loss of BRCA1 protein expression. This cell line was employed for reconstitution assays using lentivirus as in (Nacson et al., 2018). We used a virus expressing BRCA1 ΔBRCT first described in Nacson et al. (2018) and introduced the I26A mutation in both the ΔBRCT vector and a corresponding vector expressing an otherwise wild-type BRCA1. We first examined BRCA1 localization to DSB sites and found that, as expected from our studies above, the BRCA1 IR-induced foci were only abrogated when we combined the BRCT domain deletion with the I26A mutation (Figure 4a,b and Figure S4a). Similarly, only the BRCA1 I26A-ΔBRCT mutant showed strongly defective RAD51 focus formation in response to IR, in line with our previous results (Figure 4c,d). Finally, we subjected these cell lines to increasing doses of the PARP inhibitor olaparib and measured clonogenic survival. We found that the single BRCA1 I26A and ΔBRCT mutants had similar level of PARP inhibitor resistance as wild type BRCA1 (Figure 4e and Figure S4b) whereas the BRCA1 I26A-ΔBRCT mutant showed PARP inhibitor hypersensitivity that approached that of the parental or control cell line that only expressed a GFP-mCherry fusion (Figure 4e and Figure S4b). We note that in our hands, the BRCA1 ΔBRCT variant shows more activity than previously reported by Nacson et al. (2018) but this unexpected difference has no impact on our conclusions. Indeed, these results confirm that BRCA1 recruitment to DNA damage sites and its DNA repair activity involve two redundant pathways: one that involve the interaction of the BRCT domain with BRCA1-A and a second that involves the RING domain. This activity of the RING domain is also completely dependent on its interaction with its cognate E2.

**Figure 4.**
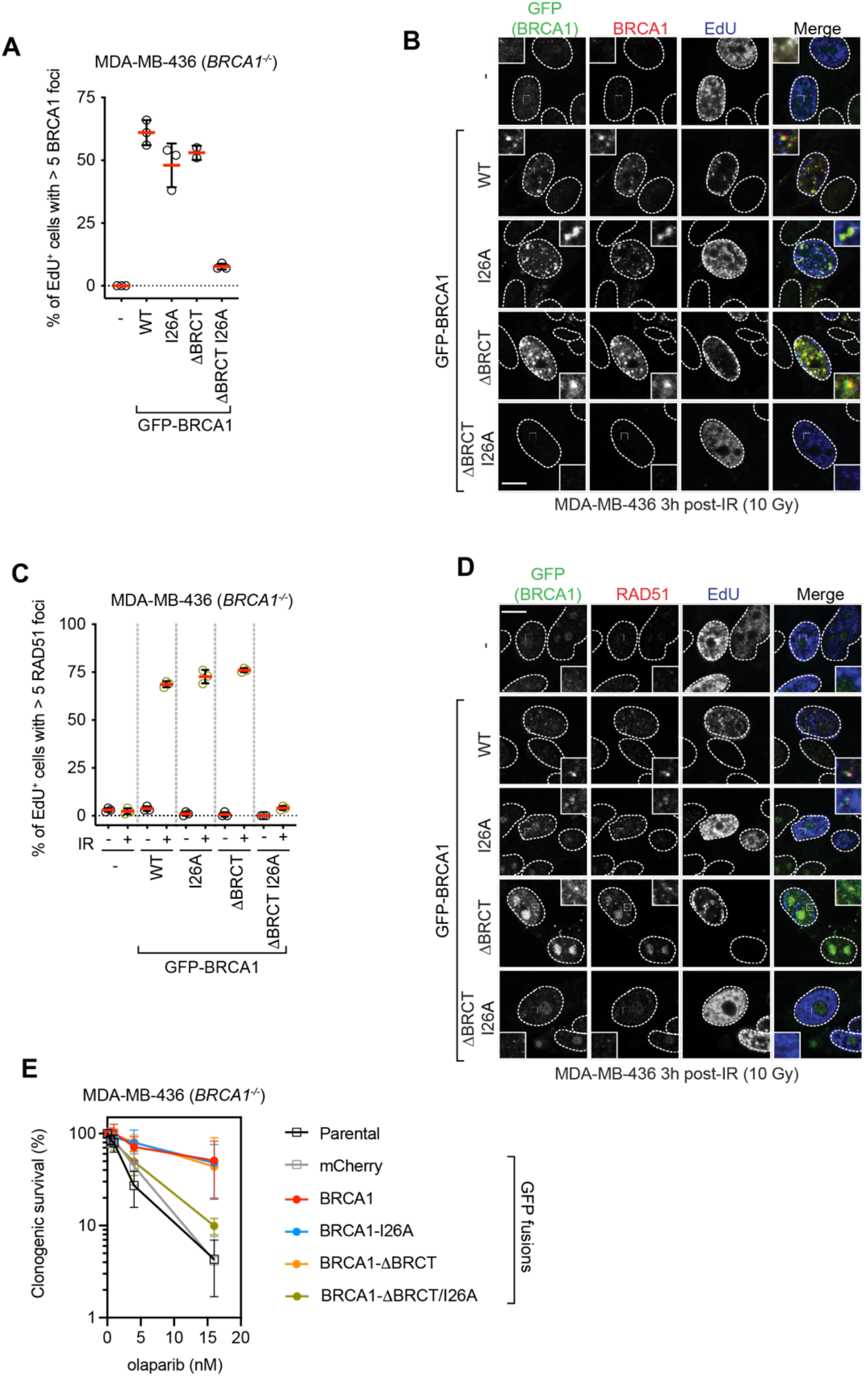
Loss of BRCA1 recruitment to DSB sites causes sensitization to PARP inhibition. (A, B) MDA-MB-436 cells transduced with lentivirus expressing the indicated GFP fusions were irradiated (10 Gy) and 3 h later were pulsed with EdU to label S phase cells 20 min prior to fixation. Cells were processed for EdU labeling and immunofluorescence using GFP and γH2AX antibodies. Shown in (A) is the quantitation of a minimum of 100 EdU^+^ cells per replicate and the bars represent the mean ± S.D (n= 3). Representative micrographs are shown in (B). (C, D). MDA-MB-436 cells transduced with lentivirus expressing the indicated GFP fusions were irradiated (10 Gy) and 3 h later were pulsed with EdU to label S phase cells 20 min prior to fixation. Cells were processed for EdU labeling and immunofluorescence using GFP and RAD51 antibodies. Shown in (C) is the quantitation of minimum of 100 EdU^+^ cells per replicate and the bars represent the mean ± S.D (n= 3). Representative micrographs are shown in (E). Clonogenic survival assays using the indicated dose or olaparib and MDA-MB-436 cells transduced with lentivirus expressing the indicated GFP fusions. Data is shown as the mean ± S.D (n=3). All scale bars are 5 μm.

## Discussion

Our results are consistent with BRCA1 having two ubiquitin-dependent modes of recruitment to DSB sites. The first recruitment pathway is dependent on the recognition of UbK63-linked chains by the RAP80 subunit of the BRCA1-A complex that interacts with BRCA1 via the latter’s BRCT domains. The second mode of recruitment involves the BRCA1 RING domain and is critical for the recognition of H2AK13/K15ub, the mark catalyzed by RNF168. These conclusions are remarkably consistent with earlier structure-function studies published over 15 years ago by Au and Henderson (Au and Henderson, 2005) and a recently posted preprint (Krais et al., 2021).

Our results suggest the BRCA1 RING domain has two distinct molecular roles that converge on the recognition of the RNF168-catalyzed histone marks at DNA damage sites. The first is to promote the interaction with BARD1. This is critical given that the H2AK13/K15ub mark is itself recognized by BARD1 via the Ankyrin repeat domain (ARD) in tandem with the BARD1 BRCT domain (the so-called BUDR) (Becker et al., 2020; Dai et al., 2021). However, the BRCA1 I26A and K70A/R71A mutations do not perturb interaction with BARD1 and therefore they must impact H2AK13/K15ub recognition by a mechanism unrelated to the formation of the BRCA1-BARD1 heterodimer. We envisage two possibilities: first, given that the I26A mutant is greatly impaired for ubiquitylation (Brzovic et al., 2006; Kalb et al., 2014), our results support a model whereby BRCA1 catalyzes C-terminal H2A ubiquitylation to promote its own recruitment. While this model is attractive, it is not clear how C-terminal H2A ubiquitylation could act to retain the BRCA1-BARD1 complex at DSB sites. An interesting idea is that the described recruitment of the SMARCAD1 chromatin remodeler by H2A K125/K127/K129 ubiquitylation displaces 53BP1 from nucleosomes (Densham et al., 2016; Densham and Morris, 2017), allowing for BRCA1- BARD1 binding to the liberated H2AK13/K15ub-bearing nucleosomes.

We also consider an alternative model where the integrity of the BRCA1 RING-E2 interaction is important for the correct positioning of the BARD1 BUDR for H2AK13/15ub recognition. Such a function for the BRCA1 RING in enabling ubiquitylated nucleosome binding is further supported with our observation that mutation of the acidic patch-interacting K70/R71 residues also impairs BRCA1 recruitment to DNA lesions in the absence of RAP80. However, we note that in a recent cryo-EM structure of the BRCA1-BARD1 complex bound to unmodified nucleosomes, the E2 is tilted away from the nucleosome and does not appear to participate in nucleosome binding (Witus et al., 2021). Furthermore, the phenotypic impact of the I26A mutation is much greater than that of the K70A/R71A mutation, which does not align as well with how these mutations impair interaction with nucleosomes and instead follows more closely how these mutations impact BRCA1/BARD1 E3 ligase activity. These two models have testable hypotheses that will be the focus of future experiments.

The observation that multiple established and likely pathogenic mutations in BRCA1 affect the RING domain (Bouwman et al., 2020; Findlay et al., 2018) and that RING-less BRCA1 protein isoforms can promote resistance to DNA damage (Drost et al., 2016; Wang et al., 2016) suggest a possible avenue for the development of agents that overcome resistance caused by RING-less BRCA1 proteins. Indeed, targeting members of the BRCA1-A complex in ways that would impair UbK63 recognition by RAP80 or that target the binding of phosphorylated ABRAXAS1 by the BRCT domains should render RING-mutated BRCA1 variants hypersensitive to PARP inhibitors or cisplatin while being relatively innocuous to cells with wild type BRCA1. While this may be difficult to accomplish through a conventional inhibitor strategy, we note that approaches such as targeted protein degradation (Neklesa et al., 2017) could provide a route for the development of agents that disrupt the DSB recruitment and HR activity of RING-less BRCA1 variants.

## Acknowledgements

We thank Cristina Escribano-Diaz for her initial observations on RAP80 and BRCA1 that helped spur this project, Rachel Szilard for critical reading of the manuscript and members of the Durocher lab for helpful discussions. We also thank Neil Johnson, Georges Mer and Ross Chapman for sharing and discussing results prior to publication. We are indebted to Roger Greenberg for the U2OS 2-6-5 cell line, Neil Johnson for sharing lentiviral expression vectors for BRCA1, Annie Bang for expert assistance with flow cytometry and Misha Bashkurov for help with image analysis. AS was supported for most of this study by a CIHR doctoral award. Work in the DD lab was supported by grants from CIHR (FDN143343) with additional support from the Krembil Foundation.

## Author Contributions

**Alana Sherker:** Conceptualization, Investigation, Writing - Original Draft, Writing - Review & Editing, Visualization; **Natasha Chaudhary:** Investigation, Resources**: Salomé Adam:** Investigation, Visualization, **Sylvie M. Noordermeer:** Resources, Review & Editing; **Amélie Fradet-Turcotte:** Resources, Review & Editing; **Daniel Durocher:** Conceptualization, Writing - Original Draft, Writing - Review & Editing, Visualization, Supervision, Funding acquisition.

## Methods

### Cell lines and culture conditions

RPE1-hTERT, HEK293 and MDA-MB-436 cells were obtained from ATCC and cultured in DMEM (Life Technologies). U2OS cells were obtained from ATCC and were cultured in McCoy’s 5A (Life Technologies). U2OS Flp-In/T-REx cells were a kind gift from Brad Wouters (OICR) and were cultured in McCoy’s 5A medium and supplemented with 15.5 μg/mL zeocin (Invitrogen) and 5 μg/mL blasticidin (Invitrogen). U2OS 2-6-3 cells (U2OS ER-mCherry-LacI-FokI-DD cells) were a kind gift from Roger Greenberg (University of Pennsylvania, Philadelphia, PA, USA) and were cultured and transfected as previously described (Shanbhag et al., 2010). All culture media was supplemented with 10% fetal bovine serum (FBS, Wisent). Cell lines were grown in an environmental incubator at 37° C and 5% CO_2_ except for BRCA1-deficient cells, which were cultured in low oxygen (3% O_2_, 5% CO_2_ at 37° C).

All cell lines were routinely authenticated by STR analysis and rested to ensure the absence of mycoplasms.

All inducible BRCA1 cell lines were generated using Flp-In/T-Rex system as described by the manufacturer (Invitrogen). Briefly, pDEST-pcDNA5-FRT/TO-GFP-derived plasmids were co-transfected with the pOG44 vector (encoding Flp recombinase) into U2OS Flp-In/T-REx cells. Recombination events were selected for with 200 μg/mL hygromycin B (Roche). Expression of BRCA1 variants was induced by the addition of 5 μg/mL doxycycline (Inalco) for 24 h. (Cell lines were obtained from ATCC (RPE hTERT, U2OS, HEK293T, MDA-MB-436) and tested for mycoplasma. Cell lines were cultured in an environmental incubator set at 37° C and 5% CO_2_ except for BRCA1-deficient cell lines that were cultured in low oxygen conditions (3% O_2_, 5% CO_2_ at 37° C). Cells were grown on plastic dishes and flasks using standard tissue culture practice. Cells were frozen in 10% DMSO in medium using Mr. Frosty containers (Nalgene) according to manufacturer’s instructions. For long term storage cells were moved to liquid nitrogen. All culture media were supplemented with 10% fetal bovine serum (FBS). U2OS cells were cultured in McCoy’s medium. HEK293T, RPE1-hTERT, MDA-MB-436 cells were cultured in DMEM.

### RNA interference

U2OS, U2OS Flp-In/T-REx, U2OS 2-6-3, RPE1-hTERT *TP53*^-/-^ *RNF8^-/-^* Cas9, or RPE1-hTERT *TP53*^-/-^ *RNF168^-/-^* Cas9 cells were transfected with siRNA using either a forward or reverse transfection mode. In both cases complexes were formed in serum-free media (Opti-MEM; Life Technologies) by adding siRNA (dissolved in 1x RNA buffer; Thermo Fisher Scientific) and the lipid-based transfection reagent RNAiMAX (Invitrogen) according to the manufacturer’s instructions. The final concentration of siRNA complexes was 10 nM. The following OnTarget Plus siRNA reagents were purchased from Horizon Discovery/Dharmacon: BRCA1 (D-003461- 05), RAP80 pool (L-006995-00-0020), BARD1 pool (L-003873-00-0020), RNF8 pool (L-006900- 00-0020), RNF168 pool (L-007152-00-0020).

### Gene-edited cell lines

Cell lines were generating using U2OS, U2OS Flp-In/T-REx, U2OS 2-6-3, or RPE1-hTERT *TP53*^- /-^ Cas9 as parental cell lines. The *RAP80*^-/-^ knockout cell lines were generated by electroporation (Lonza Amaxa II Nucleofector) of plasmids expressing sgRNAs (5′- ATTGTGATATCCGATAGTGAT-3′ and 5′-GTTCTGTCAGTGTGAAGAGG-3′) and Cas9 followed by the 2A-Puromycin cassette (pX459, Addgene #62988). 24 h after transfection, cells were selected with puromycin for 24-48 h 1 μg/mL for U2OS, U2OS Flp-In, and U2OS 2-6-5 cell lines, and 10-15 μg/mL for RPE1-hTERT *TP53*^-/-^ Cas9, and RPE1-hTERT *TP53*^-/-^ *BRCA1*^-/-^ Cas9 cell lines, followed by single clone isolation. Clones were screened by immunoblotting and immunofluorescence to verify the loss of RAP80 expression and subsequently characterized by PCR and sequencing. The genomic region targeted by the CRISPR–Cas9 was amplified by PCR using Turbo Pfu polymerase (Agilent) and the PCR product was cloned into the pCR2.1 TOPO vector (Invitrogen) before sequencing. The RPE1-hTERT *TP53*^-/-^ *BRCA1*^-/-^ Cas9 cell lines were described previously (Noordermeer et al., 2018). The RPE1-hTERT *TP53*^-/-^ *RNF8^-/-^* Cas9 cell line was generated by electroporation of phU6-gRNA (Addgene #53188) plasmids expressing sgRNAs (5’-CCCAGAGTCTAAATGGTGTT-3’ and 5’-GGAAGAGGAACAGCATCTTC-3’). Cells were isolated for single clones. The RPE1-hTERT *TP53*^-/-^ *RNF168^-/-^* Cas9 cell line was generated by electroporation of px459 plasmids expressing sgRNAs (5’-CGCTCTAAGCTTGCACTCCC- 3’ and 5’-GCCGGGTATCGTCGTGGACT-3’). Cells were selected with 15 μg/mL of puromycin, followed by single clone isolation. Both RPE1-hTERT *TP53*^-/-^ *RNF8^-/-^* Cas9 and RPE1-hTERT *TP53*^-/-^ *RNF168^-/-^* Cas9 clones were screened by immunofluorescence of BRCA1 and immunoblotting of RNF8 and RNF168 to verify the loss of RNF8 or RNF168 expression.

### Plasmids

To generate the BRCA1 expression vectors used in this study we generated Gateway-compatible entry clones in pDONR221 (Invitrogen) by amplifying BRCA1 coding regions from GFP-BRCA1 expression vectors (gift from Jiri Lukas). These entry clones were used to generate destination vectors in pDEST-pcDNA5-GFP-FRT/TO (kind gift of Anne-Claude Gingras). The deletion mutants of BRCA1 were created by deletion PCR. To generate the BRCA1 GFP-BRCA1^BRCT^ expression vector, we PCR-amplified the region encoding amino acid residues 1582-1863 from GFP-BRCA1 expression vector and ligated the product in the BamHI and EcoRI restriction sites of pcDNA5-GFP-FRT/TO. The BRCA1-I26A, BRCA1-K70A/R71A, BRCA1-S1655A, BRCA1- I26A/S1655A, and GFP-BRCT-S1655A were generated by site directed mutagenesis PCR (QuikChange; Agilent) and all plasmids were sequence verified. To generate BRCA1 constructs resistant to BRCA1 siRNA #5 (Dharmacon, D-003461-05), we introduced the following underlined silent mutations in BRCA1: 5’<u>AGT</u>AT<u>A</u>AT<u>C</u> 3’. For CRISPR-Cas9 genome editing, DNA corresponding to sgRNAs was cloned into pX459 (Addgene #62988). The pDEST-IRES-GFP vectors containing BRCA1, BRCA1-ΔBRCT were constructed and provided by Neil Johnson (Nacson et al., 2018). The BRCA1-I26A mutation was generated by site directed mutagenesis (Agilent) using the BRCA1 and BRCA1-ΔBRCT in the pENTR1A Gateway Entry vector (Invitrogen) and shuttled into a pDEST-IRES-GFP Destination vector (Life Technologies) using Gateway LR Clonase II Enzyme Mix (Invitrogen).

### Immunofluorescence microscopy

Cells were grown on glass coverslips, ionizing radiation was delivered with a Faxitron X-ray cabinet (#43855D) at 120 kV output voltage, 3 mA continuous current, 3.07 Gy/min. Unless otherwise stated, 1 h post-IR cells were fixed with 4% (w/v) paraformaldehyde in PBS for 10 min at room temperature, permeabilized with 0.25% (v/v) Triton X-100 + 2.5% BSA PBS for 10 min at room temperature and blocked with PBS + 0.1% Tween-20 + 5% BSA PBS for 30 min at room temperature. Alternatively, cells were pre-extracted 10 min on ice with NuEx buffer (20 mM HEPES, pH 7.4, 20 mM NaCl, 5 mM MgCl2, 0.5% NP-40, 1 mM DTT and a cocktail of protease inhibitors (cOmplete, EDTA-free; Roche)) followed by 10 min 4% PFA fixation. Cells were then incubated with the primary antibody diluted in PBS + 0.1% Tween-20 + 5% BSA for 1-2 h at room temperature. Cells were next washed with PBS and then incubated with secondary antibodies diluted in PBS + 5% BSA supplemented with 0.8 μg/mL of DAPI to stain DNA for 1 h at room temperature. Secondary antibodies were purchased from Molecular Probes (Alexa Fluor 488 goat anti-mouse IgG, Alexa Fluor 488 goat anti-rabbit IgG, Alexa Fluor 488 donkey anti-mouse IgG, Alexa Fluor 55 goat anti-mouse IgG, Alexa Fluor 555 goat anti-rabbit IgG, Alexa Fluor 647 goat anti-mouse IgG, Alexa Fluor 647 goat anti-rabbit IgG). The coverslips were mounted onto glass slides with Prolong Gold mounting agent (Invitrogen). Confocal images were taken using a Zeiss LSM780 laser-scanning microscope. For examination of S-phase, cells were pre-incubated with 20 μM of EdU (Life Technologies; A10044) for 30 min after irradiation and 30 min prior to fixation and processed as follows: Cells were incubated with primary antibody diluted in PBS + 0.1% Tween-20 + 5% BSA for 1 h at room temperature. Cells were next washed with PBS and then incubated with secondary antibodies diluted in PBS + 5% BSA supplemented with 0.8 μg/mL of DAPI to stain DNA for 1 h at room temperature. Cells were then fixed for 5 min in 4% PFA, washed and stained with EdU staining buffer (100mM Tris-HCl pH 8.5, 10 μM Alexa Fluor 647, 1 mM CuSO_4_, 100 mM ascorbic acid). Unless otherwise stated analysis was performed on 100 cells per condition, where cells with greater than 10 (GFP, BRCA1) or 5 (RAD51) irradiation-induced foci that co-localize with γH2AX were considered positive. Experiments were performed in triplicates. For experiments in U2OS-2-6-3 cells, 25-30 images per condition were acquired and GFP intensity at the mCherry-FokI focus was measured relative to the background intensity of the nucleus. To differentiate the true signal present in BRCA1 (WT) and RING (BRCA1 1-110) samples from noise, a clustering mixture model was used, assuming the GFP control would follow a Gaussian distribution and the true signal would follow a gamma distribution. First, the Gaussian parameters were estimated by the method of moments on the GFP control data alone; second, for the other samples, these parameters were held static while the gamma parameters were also estimated using the method of moments and optimized through the expectation maximization algorithm. The data shown in Figure S2a shows the GFP control signal and the true signal estimated in the GFP-BRCA1 and -RING samples (i.e. with noise subtracted).

### Lentiviral production and cell line generation

HEK293T cells were transfected with pDEST-IRES-GFP BRCA1 variant plasmids along with psPAX2 packaging and VSV-G/pMD2.G envelope plasmids as described previously (Nacson et al., 2018). MDA-MB-436 cells were infected with the resulting lentivirus as described previously (Nacson et al., 2018) and infected cells were selected based on their GFP expression, where cells were double-sorted for populations expressing GFP-mCherry, -BRCA1, -BRCA1-I26A, -BRCA1- ΔBRCT, and -BRCA-ΔBRCT-I26A. Cells were tested for protein expression by immunoblotting.

### Clonogenic survival assays

Cells were seeded in 10-cm dishes (500-1000 cells per 10 cm plate, depending on cell line and genotype). For drug sensitivity assays cells were seeded into media containing a range of olaparib (SelleckChem S1060) concentrations. Plates were cultured in low oxygen conditions (3% O_2_, 5% CO_2_ at 37° C). Medium was refreshed after 7 days in all cases. At the end of the experiment medium was removed, cells were rinsed with PBS and stained with 0.4% (w/v) crystal violet (Sigma; C0775) in 20% (v/v) methanol for 30 min. The stain was aspirated, and plates were rinsed twice in deionized H_2_O and air-dried. Colonies were counted using a GelCount (Oxford Optronix) and data were plotted as surviving fractions relative to untreated cells.

### Antibodies

We employed the following antibodies for immunofluorescence: mouse anti-γH2AX (clone JBW301, Millipore, 1:1000), mouse anti-BRCA1 (clones MS110 and MS13 Calbiochem, 1:100), rabbit anti-BRCA1 (#07-434, Millipore 1:1000), rabbit anti-RAD51 serum (70-001; lot 1; BioAcademia, 1:15,0000). Rabbit anti-RAP80 Rabbit (A300-763A, Bethyl Laboratories Inc., 1:200) anti-RAP80 Rabbit (NBP1-87156, Novus Biologicals, 1:500) anti-ABRAXAS (A302- 180A, Bethyl Laboratories Inc., 1:500) We employed the following antibodies for immunoblotting: rabbit anti-BRCA1 (homemade, 1:1000) (Noordermeer et al., 2018), rabbit anti- RNF8 (kind gift from Junjie Chen, 1:2500), rabbit anti-RNF168 (homemade, 1:2500) (Stewart et al., 2009) , rabbit anti-RAP80 (A300-763A, Bethyl Laboratories Inc., 1:5000), mouse anti-tubulin (clone DM1A, Calbiochem, 1:1000), rabbit anti-KAP1 (Bethyl, 1:10,000). Rabbit anti-GAPDH (G9545, Sigma Aldrich, 1:5,000).

### Traffic light reporter assay

RPE1-hTERT *TP53*^-/-^ Cas9 and RPE1-hTERT *TP53*^-/-^ *RAP80*^-/-^ Cas9 cells were transduced with pCVL.TrafficLightReporter.Ef1a.Puro (Noordermeer et al., 2018) lentivirus at a low MOI (0.3) and selected with puromycin (15 μg/mL). Cells (7×10^5^) were nucleofected with 5 μg of pCVL.SFFV.d14mClover.Ef1a.HA.NLS.Sce(opt).T2A.TagBFP (Noordermeer et al., 2018) plasmid DNA in 100 μL electroporation buffer (25 mM Na_2_HPO_4_ pH 7.75, 2.5 mM KCl, 11 mM MgCl_2_), using program T23 on a Nucleofector 2b (Lonza). After 48-72 h, mClover fluorescence was assessed in BFP-positive cells using a Fortessa X-20 (BD Biosciences, San Jose) flow cytometer.

---

**Figure S1.**
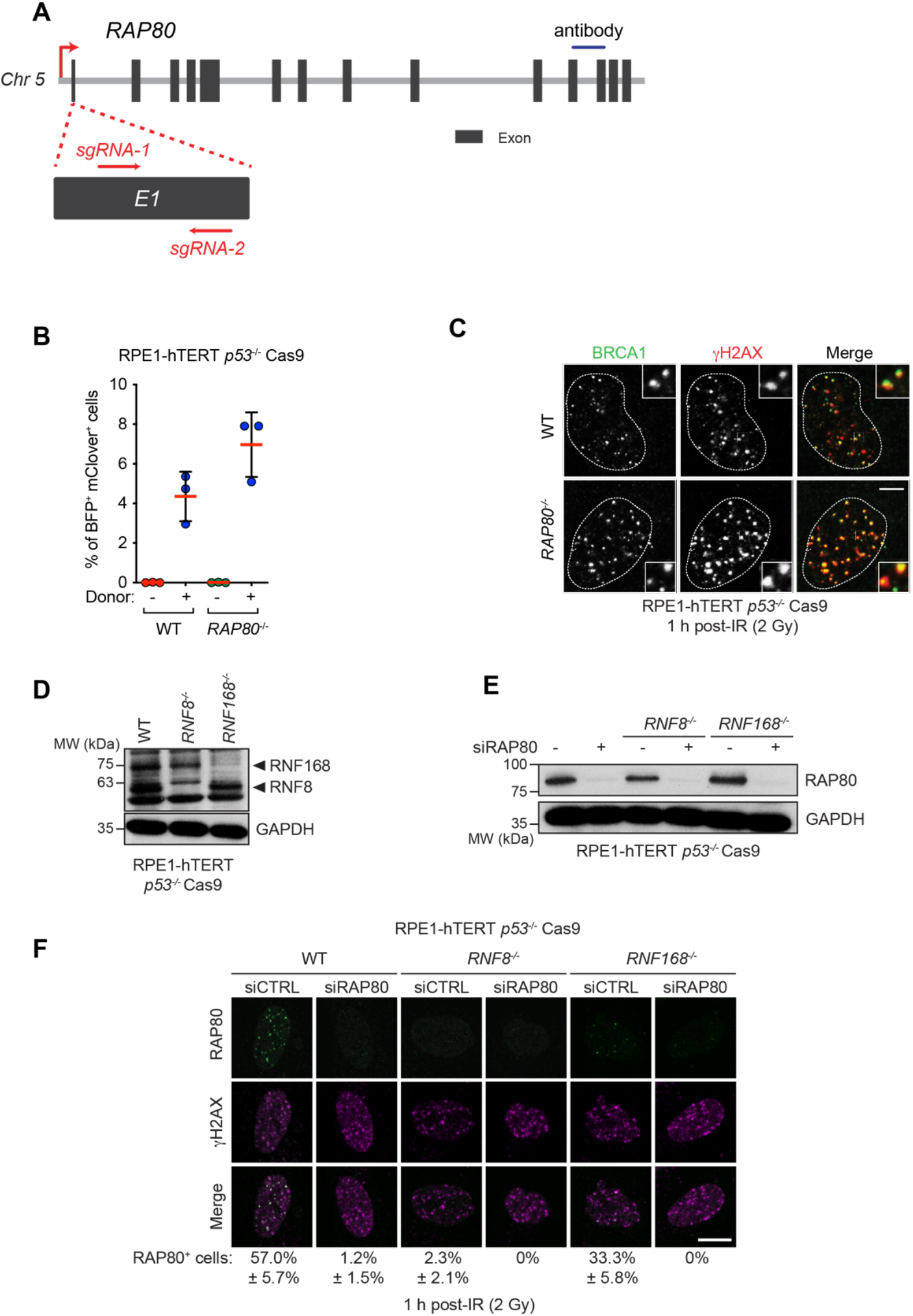
Design and characterization of *RAP80*^-/-^ cell lines. (A) To generate *RAP80^-/-^* cell lines, we designed single guide RNAs (sgRNA) that target the first exon of *RAP80*. U2OS and U2OS Flp-In/T-REx clones were sequence validated (Table 1) and carry RAP80 indels leading to premature stop codons. Dark grey boxes indicate exons 1-14, red arrows indicate the location of the sgRNA, blue line indicates the epitope recognized by the RAP80 used for immunoblotting. (B) Homologous recombination was monitored by gene conversion using the traffic light reporter assay in the indicated cell lines 48 h post-nucleofection in the absence or presence of the donor template. BFP^+^ mClover^+^ cells have undergone gene conversion. The bars represent the mean ± S.D (n=3). (C) Representative micrographs of the experiment shown in Figure 1g at the 1 h timepoint. (D) Immunoblotting of whole cell extracts obtained from parental (WT), *RNF8^-/-^*, and *RNF168^-/-^* RPE1-hTERT *p53^-/-^* Cas9 cells with a mixture of RNF8 and RNF168 antibodies. GAPDH is used as a loading control. (E) Immunoblotting of whole cell extracts from the indicated cell lines treated with either a non-targeting siRNA control (CTRL) or siRNA targeting *RAP80.* GAPDH is used as a loading control. (F) The indicated parental (WT), *RNF8^-/-^*, and *RNF168^-/-^* cell lines were processed for immunofluorescence 1 h post-irradiation (2 Gy) and stained with antibodies against RAP80 and γH2AX. Representative micrographs are shown with quantitation of the mean percentage of cells with > 5 RAP80 foci that colocalize with γH2AX written below (n=3). A minimum of 50 cells were analyzed. All scale bars are 10 μm.

**Figure S2.**
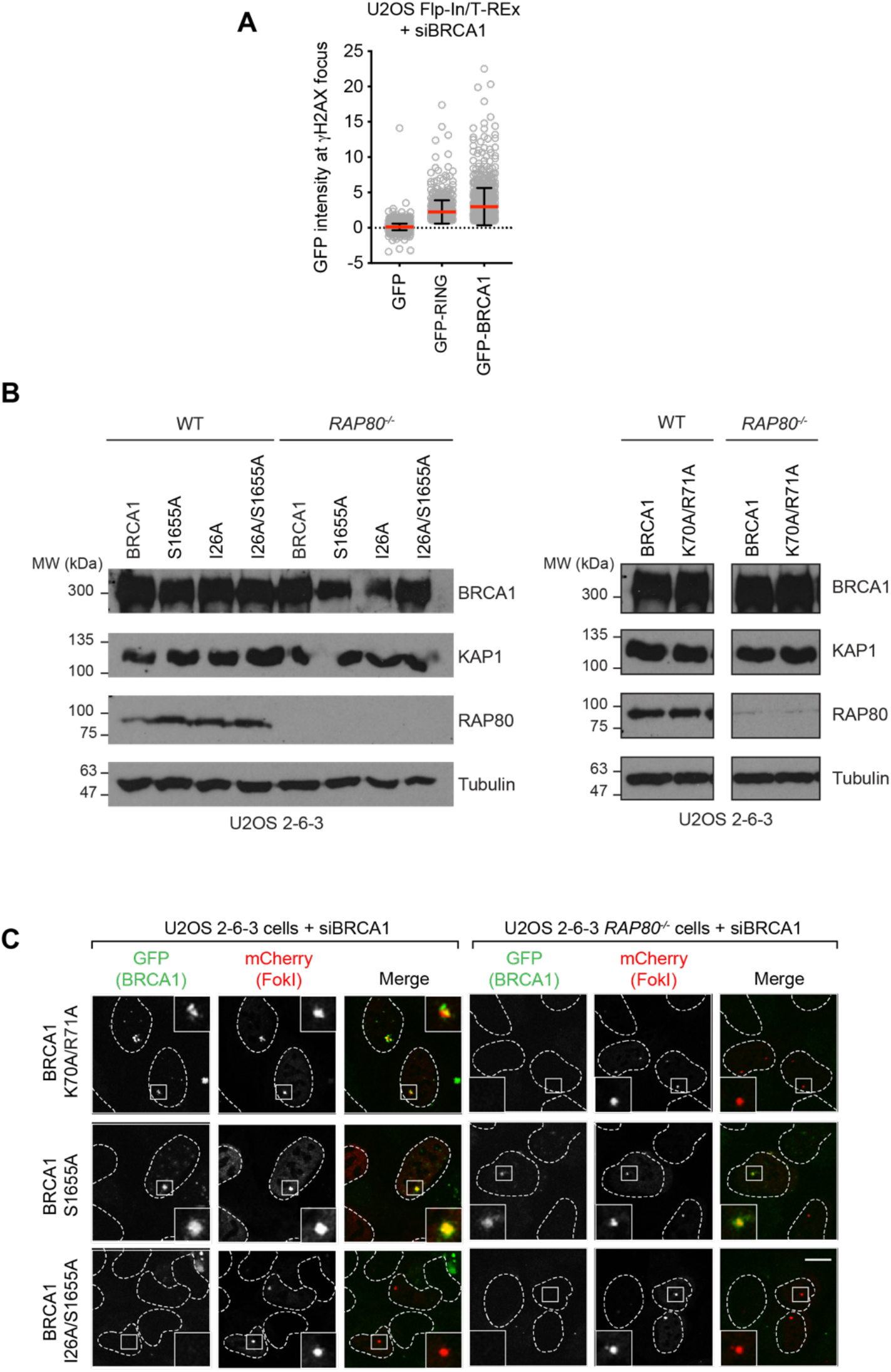
Supplementary information for Figure 3. (A) Quantitation of GFP intensity at the γH2AX focus in U2OS Flp-In/T-REx cells expressing the indicated GFP fusion proteins. An Acapella analysis was performed: γH2AX IR-induced foci were found within a nucleus and the GFP intensity under each γH2AX focal unit was measured. Intensity of GFP under each individual γH2AX focus was plotted with the background GFP intensity subtracted. The red line indicates the mean for each condition. (B) Immunoblotting of parental U2OS 2-6-3 (WT) or isogenic *RAP80^-/-^* cell lines transfected with siRNAs targeting *BRCA1* and nucleofected with vectors expressing the indicated GFP fusion proteins as described in Figure 3c,d. Tubulin and KAP1 are used as loading controls. (C) Representative micrographs for the additional conditions shown in Figure 3c. All scale bars are 10 μm.

**Figure S3.**
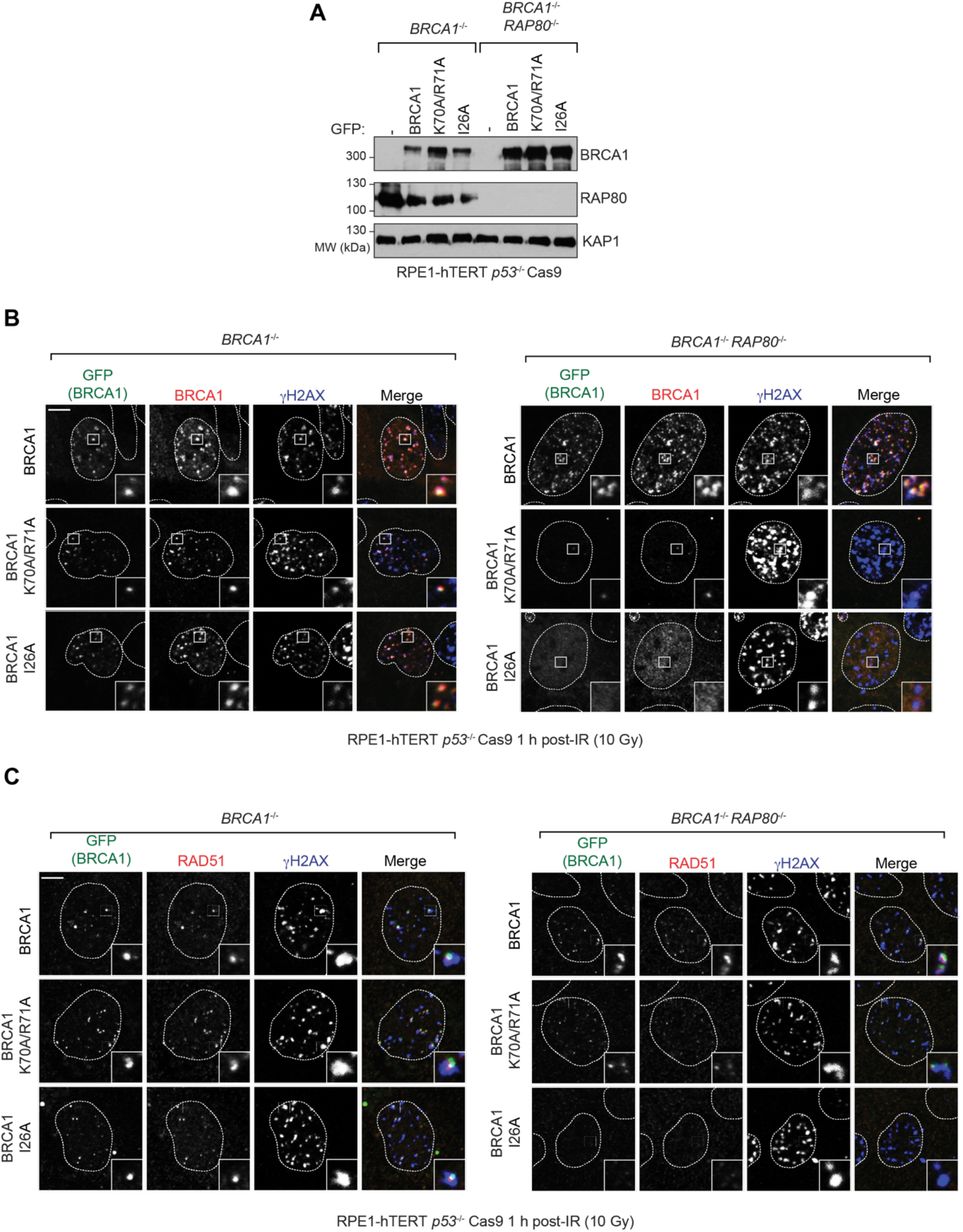
Additional controls for Figure 3. (A) Whole cell extracts from RPE-1 hTERT *p53^-/-^ BRCA1^-/-^* Cas9 cells or their isogenic *RAP80^-/-^* counterparts expressing the indicated GFP fusion proteins were immunoblotted with antibodies against BRCA1, RAP80 and KAP1 (loading control). Relates to Figure 3e,f (B,C) RPE-1 hTERT *p53^-/-^ BRCA1^-/-^* Cas9 cells or their isogenic *RAP80^-/-^* expressing the indicated GFP fusion proteins were processed 1 h post-irradiation (10 Gy) for immunofluorescence using GFP and antibodies against γH2AX, and either BRCA1 (B) or RAD51 (C). Relates to Figure 3e,f. Scale bars are 5 μm.

**Figure S4.**
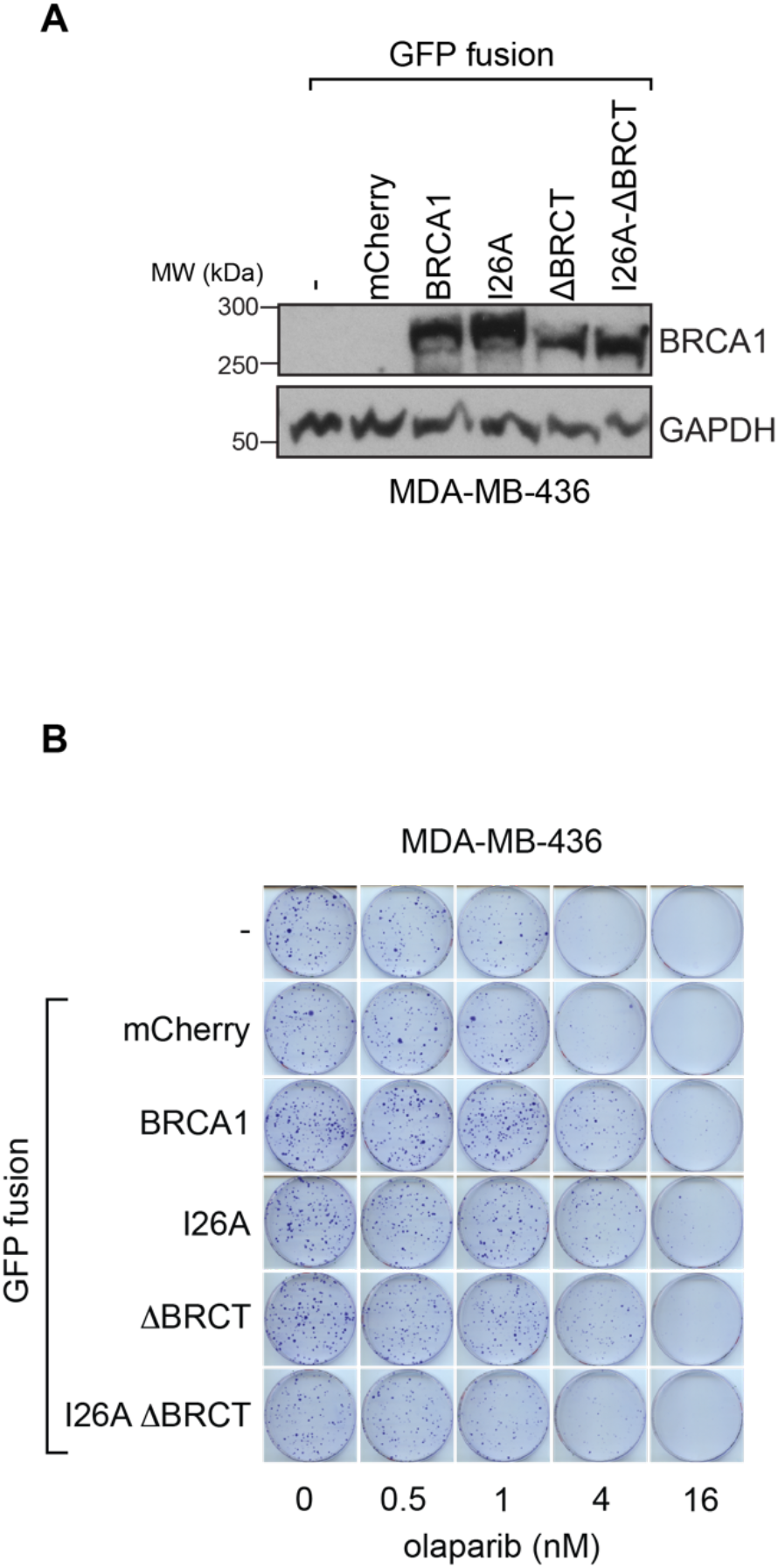
Additional controls and information for Figure 4. (A) Immunoblotting of whole cell extracts were obtained from MDA-MB-436 cells transduced with lentivirus expressing the indicated GFP fusions. GAPDH was used as a loading control. (B) Representative images of clonogenic survival assays using the indicated dose of olaparib and MDA-MB 436 cells transduced with lentivirus expressing the indicated GFP fusions.

